# DnoisE: Distance denoising by Entropy. An open-source parallelizable alternative for denoising sequence datasets

**DOI:** 10.1101/2021.07.07.451520

**Authors:** Adrià Antich, Creu Palacín, Xavier Turon, Owen S. Wangensteen

## Abstract

DNA metabarcoding is broadly used in biodiversity studies encompassing a wide range of organisms. Erroneous amplicons are generated during amplification and sequencing procedures and constitute one of the major sources of concern for the interpretation of metabarcoding results. Several denoising programs have been implemented to detect and eliminate these errors. However, almost all denoising software currently available has been designed to process non-coding ribosomal sequences, most notably prokaryotic 16S rDNA. The growing number of metabarcoding studies using coding markers such as COI or RuBisCO demands a re-assessment and calibration of denoising algorithms. Here we present DnoisE, the first denoising program designed to detect erroneous reads and merge them with the correct ones using information from the natural variability (entropy) associated to each codon position in coding barcodes. We have developed an open-source software using a modified version of the UNOISE3 algorithm. DnoisE implements different merging procedures as options, and can incorporate codon entropy information either retrieved from the data or supplied by the user. In addition, the algorithm of DnoisE is parallelizable, greatly reducing run times on computer clusters. Our program also allows different input file formats, so it can be readily incorporated into existing metabarcoding pipelines.

## Background

Biodiversity studies have experienced a revolution in the last decade with the application of high throughput sequencing (HTS) techniques. In particular, the use of metabarcoding in ecological studies has increased notably in recent years. For both prokaryotic and eukaryotic organisms, a large number of applications have been developed, ranging from biodiversity assessment, detection of particular species (Kelly et al., 2014), analysis of impacts (Pawlowski et al., 2018), and diet studies (Clarke et al., 2018; Sousa et al., 2019), among others. Also, different sample types have been used: terrestrial soil, freshwater, marine water, benthic samples, arthropod traps, or animal faeces (Creer et al. 2016, Deiner et al. 2017). Many of these studies have direct implications on management and conservation of ecosystems and are thus providing direct benefits to society. They have also brought to light a bewildering diversity of organisms in habitats difficult to study with traditional techniques.

Metabarcoding studies have greatly contributed to so-called big community data (Pichler & Hartig 2020), by generating an enormous amount of sequence data that, in most cases, is available online. Handling these datasets is memory intensive and filtering steps are required to analyze such information. Clustering and denoising are the two main strategies to compress data into Molecular Operational Taxonomic Units (MOTUs, aka OTUs) or Exact Sequence Variants (ESVs; also ASVs, Amplicon Sequence Variants, or ZOTUs, zero ratio OTUs) to extract biodiversity composition (Antich et al., 2021). Both methods rely on minimizing sequencing and PCR errors either by clustering sequences into purportedly meaningful biological entities (MOTUs) or by merging erroneous sequences with the correct ones from which they possibly originated, and keeping just correct amplicons (ESVs). Hence, both methods differ philosophically and analytically. Furthermore, they are not incompatible and can be jointly applied. Software development is crucial to create tools capable of performing these tasks in a fast and efficient way. The type of samples, the marker, and the target organisms are also instrumental in choosing the adequate bioinformatic pipelines to provide interpretable results.

Recent studies have explored the application of both methods to filter metabarcoding data (Antich et al., 2021; Brandt et al., 2021; Elbrecht et al., 2018; Turon et al., 2020). Importantly, the combination of clustering and denoising opens the door to the analysis of intraspecies (intra-MOTU) variability (Antich et al., 2021). Turon et al. (2020) proposed the term metaphylogeography for the study of population genetics using metabarcoding data, and Zizka et al. (2020) found different haplotype composition between perturbed and unperturbed rivers, both studies using a combination of clustering and denoising steps.

The software presented here focuses on the denoising step. There are currently several software programs developed to denoise sequencing and PCR errors, such as DADA2 (Callahan et al., 2016), AmpliCL (Peng & Dorman, 2020), Deblur (Amir et al., 2017), or UNOISE3 (Edgar, 2016). These programs have been widely used in metabarcoding studies to generate ESVs, using sequence quality information for the first two and simple analytical methods for the latter two. All were originally tested for ribosomal DNA (non-coding) and thus some adjustment is necessary for application to other markers (Antich et al., 2021).

Here we present DnoisE, a parallelizable Python3 software for denoising sequences using a modification of the UNOISE3 algorithm and tested for metabarcoding of eukaryote communities using mitochondrial markers (COI, Cytochrome Oxidase subunit I). We introduce a novel correction procedure for coding sequences using changes in diversity values per codon position. In coding genes, the natural entropy of the different positions is markedly different, with the third position being always the most variable. We therefore contend that differences in each position should have different weights when deciding whether a change in a given position is legitimate or is attributable to random PCR or sequencing errors. DnoisE is also applicable to other markers due to the settable options and offers a fast and open source alternative to non-parallelizable closed source programs.

## Workflow

### Structure of input files

DnoisE is designed to run with HTS datasets (after paired-end merging and de-replicating sequences) to obtain ESVs, or after clustering with SWARM (Mahé et al., 2015) to obtain haplotypes within MOTUs. Due to variability in format files, we have designed an algorithm that can read both fasta and csv files. In the present version, however, sample information (if present) is kept only for csv input.

### Combining the UNOISE3 algorithm and the entropy correction

Sequences are stored as a data frame, with each row corresponding to a sequence record and the columns to the abundances (either total or per sample). The original Edgar’s (2016) function used by UNOISE3 to determine whether two sequences should be merged is:

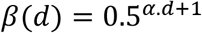

where β(d) is the threshold abundance ratio of a less abundant sequence with respect to a more abundant one (from which it differs by d) below which they are merged. The distance d is the Levenshtein genetic distance measured with the Levenshtein-python module (https://github.com/ztane/python-Levenshtein/) and α is the stringency parameter (the higher α, the lower the abundance skew required for merging two sequences).

UNOISE3 algorithm sorts sequences by decreasing abundance and each one is compared with the less abundant ones. At each comparison, the difference between sequences (d) is computed and, if the abundance ratio between the less abundant and the more abundant sequence is lower than β(d), the former is assumed to be an error. In UNOISE3 terminology, the sequences form clusters, of which the correct one is the centroid and the remaining members are inferred to derive from the centroid template but containing errors. In his original paper, Edgar (2016) suggests constructing a table of centroids excluding low abundance reads, and then constructing a ZOTU table by mapping all reads (before the abundance filtering) to the centroids table using the same merging criteria but without creating new centroids. This standard procedure of UNOISE3 gives priority to the abundance ratio over the genetic distance. The first, very abundant, sequences will “capture” rare sequences even if d is relatively high. Other, less abundant sequences may be closer (lower d) and still fulfil Edgar’s formula for merging the rare sequence, but this will never happen as the rare sequence will be joined with the very abundant one and will not be available for further comparisons. However, in the standard procedure implemented in the USEARCH pipeline (Edgar 2010, https://drive5.com/usearch/), the reads are mapped to the centroid table using a similarity criterion (identity threshold in the otutab command).

DnoisE is a one pass algorithm, with no posterior mapping of reads to centroids (which is indeed repetitive, as most reads have already been evaluated against the centroids when constructing the centroid table). If deemed necessary, low abundance reads can be eliminated previously or, alternatively, ESVs with one or a few reads can be discarded after denoising. Chimeric amplicons can likewise be eliminated before or after denoising. DnoisE follows previously used terminology (Turon et al. 2020, Antich et al. 2021) in which the correct sequences (centroids in UNOISE3 terms) are called “mother” sequences and the erroneous sequences derived from them are labelled “daughter” sequences. DnoisE provides different options for merging the sequences. Let PMS (potential “mother” sequence) and PDS (potential “daughter” sequence) denote the more abundant and the less abundant sequences that are being compared, respectively, and let d be the genetic distance between them. When the abundance ratio PDS/PMS is lower than β(d), the PDS is tagged as an error sequence but is not merged with the PMS. Instead, a round with all comparisons is performed and, for a given PDS, all PMS fulfilling the UNOISE3 criterion for merging are stored. After this round is completed, the merging is performed following three different criteria: (1) Ratio criterion, joining a PDS to its more abundant PMS (lowest abundance ratio, corresponding to the original UNOISE3 formulation); (2) Distance criterion, joining a sequence to the closest (least d value) possible “mother”; and the (3) Ratio-Distance criterion, whereby a PDS is merged with the PMS for which the quotient β/β(d) (i.e., between the abundance ratio PDS/PMS and the maximal abundance ratio allowed for the observed d), is lowest, thus combining the two previous criteria. For each criterion, the best PMS and the corresponding values (ratio, d and ratio skew values) are stored. The user has then the choice to select one or another for merging sequences. As an option, if the user wants to apply only the Ratio criterion, each PDS is assigned to the first (i.e., the most abundant) PMS that fulfils the merging inequality and becomes unavailable for further comparisons, thus decreasing computing time. Figure 1 shows a conceptual scheme of the workflow process.

**Figure 1.**
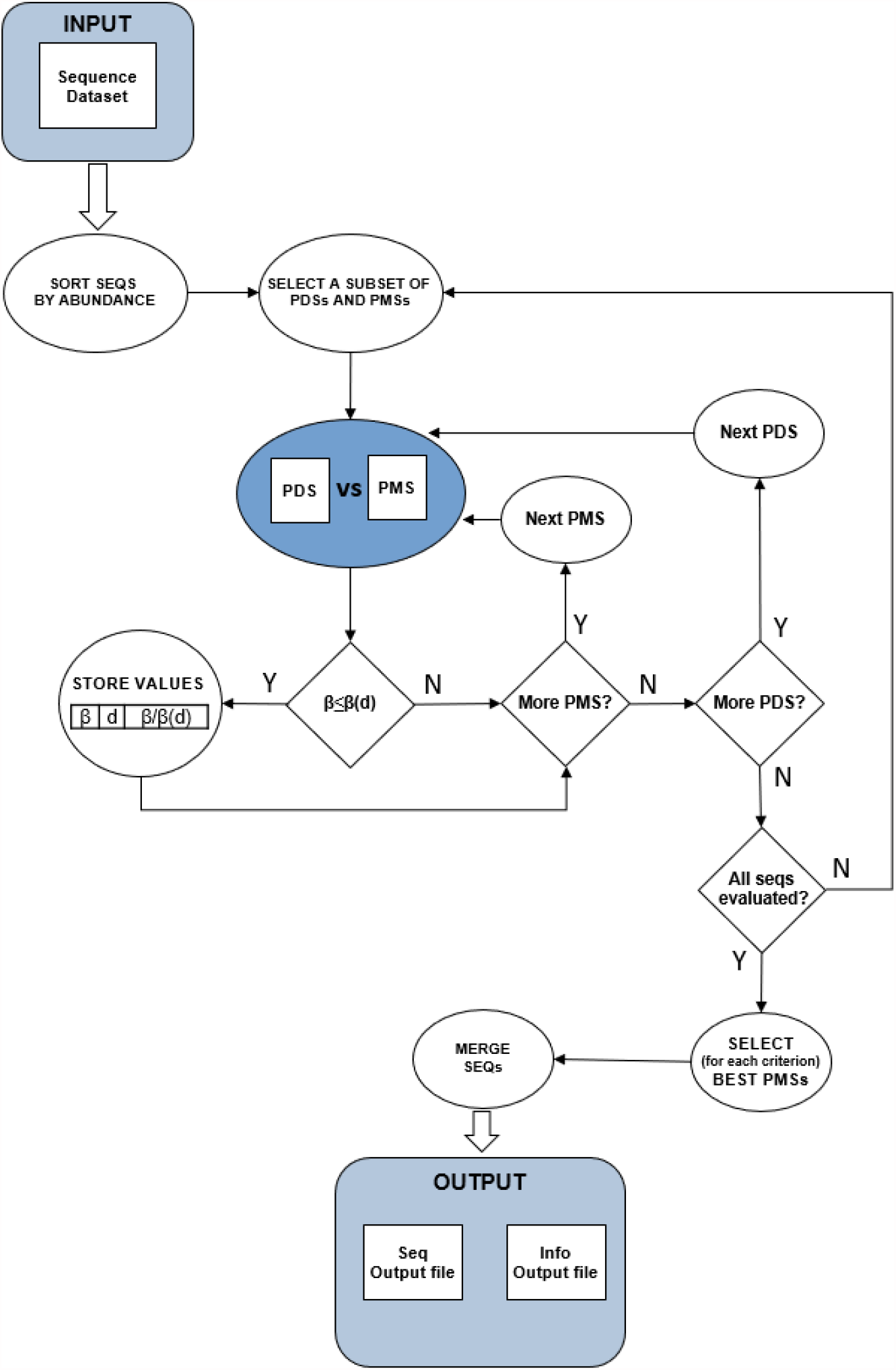
Scheme of the workflow of DnoisE. Starting from an abundance-sorted sequence dataset, subsets of PDSs and PMSs are selected as detailed in Fig. 2. For each subset, all PDSs are compared with all compatible PMSs (in terms of MDA and MMA). If the merging inequality is realized, the values of the main parameters are stored. After all subsets have been evaluated, for each merging criterion the best PMS for each PDS is chosen and a sequence file is generated, together with a file with information on the merging process.

In addition, for coding markers such as COI, the codon position provides crucial additional information that must be taken into account. In nature, the third codon position is the most variable, followed by the first and the second position. This variation can be measured as entropy (Schmidt & Herzel 1997) of the different positions. A change in third position is more likely to be a natural change (and not an error) than the same change in a second position, much less variable naturally. To our knowledge, no denoising algorithm incorporates this important information. We propose to use the entropy values of each codon position to correct the distance d in Edgar’s formula as follows:

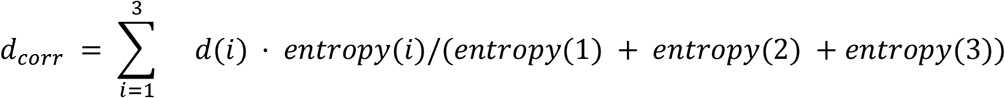

where *i* is the codon position and d is the number of differences in each position. The *d*_*corr*_ value is then used instead of *d* in the formula. This correction results in a higher *d*_*corr*_ when a change occurs in a third position than in the first or second position, thus a sequence with changes in third positions will be less likely to be merged. In practice, as many changes occur naturally in third positions, this correction will lead to a higher number of ESVs retained that would otherwise be considered errors. Careful choice of entropy values is crucial, and it is recommended that they are adjusted for each marker and particular study. The values of entropy for each position can be obtained from the data (the *entrpy*.*R* script returns entropy values) or added manually by the user. Note that, when applying this correction, the Levenshtein distance is not used as it cannot consider codon positions. Instead, the number of differences is used. In practice, in aligned sequences with no indels both distances are equivalent.

### Parallel processing

Parallel processing is a useful tool to increase speed when multicore computers are available. DnoisE implements parallel processing in the algorithm so the required time to run huge datasets decreases drastically as more cores are used. Parallel processing was applied using the multiprocessing module of Python3 (McKerns et al., 2011). A computational bottleneck of denoising procedures is their sequential nature, that is hardly parallelizable, and more so in the case of DnoisE that computes all comparisons before merging. In particular, a sequence that has been tagged as “daughter” (error) cannot be a “mother” of a less abundant sequence. Therefore, to compare a PDS to all its PMS requires that those more abundant sequences have been identified as correct before.

We incorporate two concepts, based on the highest skew ratio required for a sequence to be merged with a more abundant one. This is of course β(min(d)), where min(d) is one if entropy correction is not performed, and it equals the *d*_*cor*r_ corresponding to a single change in the position with less entropy (position 2) if entropy is considered. From this maximal abundance ratio we can obtain, for a given potential “mother”, the maximal “daughter” abundance (MDA, any sequence more abundant than that cannot be a “daughter” of the former). Conversely, for a given “daughter” sequence we can obtain the minimum “mother” abundance (MMS, any sequence less abundant than that cannot be the “mother” of the former). The formulae are:

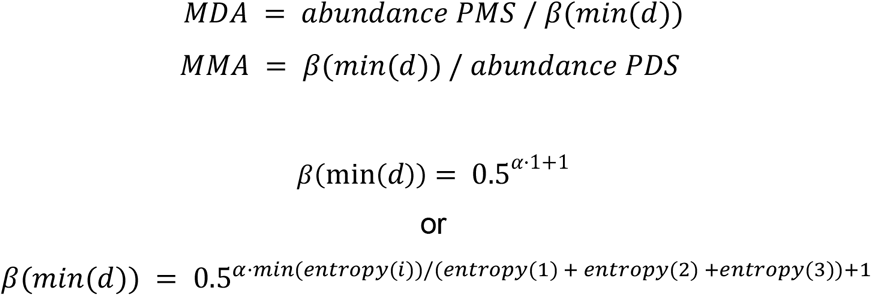

The use of MDA and MMA simplifies the workload of the program as it greatly reduces the number of comparisons (a PMS will not be evaluated against sequences more abundant than the MDA, and a PDS will not be compared with sequences with less abundance than the MMA). Likewise, it allows for a parallel processing of sequences using the MDA as follows:

1. Sequences are ordered by decreasing abundance.
2. The first sequence is automatically tagged as a correct sequence.
3. MDA is calculated for this sequence (MDA_1).
4. All sequences with abundances between the first sequence and the MDA are, by definition, tagged also as correct sequences.
5. For the last sequence tagged as correct, the MDA is calculated (MDA_2).
6. Every sequence with abundance between the last correct sequence and MDA_2 is evaluated in parallel against all correct sequences that are more abundant than its MMA. Those for which no valid “mother” is found are tagged as correct, the rest are “daughter” (error) sequences.
7. Repeat steps 5 and 6 (i.e., calculating MDA_3 to n) until all sequences have been evaluated.

Figure 2 provides a conceptual scheme of this procedure. Note that, for each block of sequences that is evaluated in parallel, no comparisons need to be performed between them as they will never fulfill the merging inequality. After this process is completed, all sequences not labelled as “daughter” are kept as ESVs, and all “daughters” are merged to them according to the merging criterion chosen.

**Figure 2.**
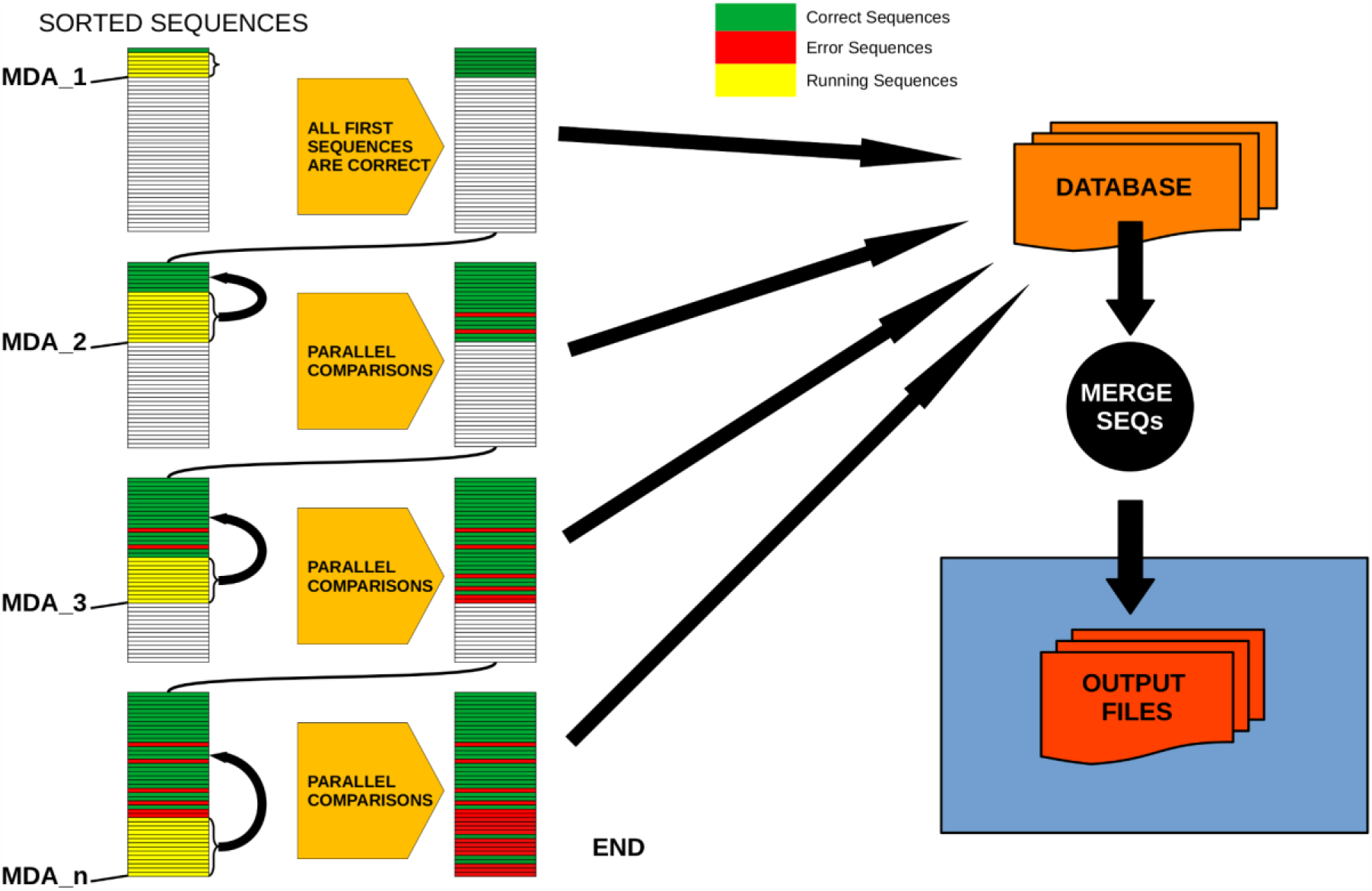
Schematic workflow of parallel processing of DnoisE. When running in parallel, comparisons between sequences are computed in sets of sequences defined by their abundances. Using the Maximum Daughter Abundance (MDA) value, computed from the last correct sequence of the previous step, we can define sets of sequences that are compared in parallel with the previously tagged correct sequences.

## DnoisE performance

A beta version of DnoisE was tested in Antich et al. (2021) on a COI metabarcoding dataset of marine benthic communities. The dataset consisted of 330,382 chimera-filtered COI sequences of 313 bp (all sequences had more than one read). They came from benthic marine communities in 12 locations of the Iberian Mediterranean coast (see Antich et al. 2021 for details), and are available as a Mendeley Dataset (https://data.mendeley.com/datasets/84zypvmn2b/). DnoisE was used in Antich et al (2021) in combination with the clustering algorithm SWARM, and was compared with the results of DADA2 denoising algorithm. Antich et al. (2021) also compared DnoisE with and without entropy correction, and obtained twice the number of ESVs with correction, while the proportion of erroneous sequences (defined as those having stop codons or substitutions in conserved positions) decreased to one half as compared with not correcting for codon position variation, as discussed in Antich et al (2021).

In the present paper, we benchmarked the current version of DnoisE (with alpha=5) against UNOISE3 (USEARCH 32-bit, free version, with alpha=5 and minsize=2) on this same dataset. The number of ESVs obtained was similar: 60,198 and 60,205, respectively, if no entropy correction was performed. In addition, 60,196 ESVs were shared (comprising >99.999% of the total reads) among the two programs.

We also compared the run speed of DnoisE with and without entropy correction for the same dataset of sequences. We used different numbers of cores, from 1 to 59, for parallelization. We applied the entropy correction values from Antich et al. (2021) (Scripts for installation and examples of running DnoisE are provided in Supplementary materials).

Running DnoisE with just one core (without entropy correction) took about 29 hours, decreasing sharply when using parallel processing with just a few cores. DnoisE took 4.5 hours with 6 cores and 2.78 hours with 10 cores. As a reference, the execution time of UNOISE3 (32-bit version, not parallelizable) without the otutab step was ca. 7 hours, albeit this execution time is not directly comparable as UNOISE3 has a chimera filtering step embedded. Using entropy correction, run times increased (Fig. 3) as there is a higher number of comparisons needed because the MMA values are in general lower. This slows the process as any given PSD has more PMS to compare with. With entropy correction, DnoisE retrieved ca. twice the number of ESVs, further increasing run time. For the Ratio-Distance merging criterion, when entropy correction was performed, 16 cores were required for DnoisE to run at a similar time speed than 6 cores with no entropy correction (Fig. 3). Above 10 cores (without correction) or 20 cores (with correction), run times reached a plateau and did not further improve, while memory usage continued to increase steadily. A trade-off between both parameters should be sought depending on the cluster architecture and the dataset being run

**Figure 3.**
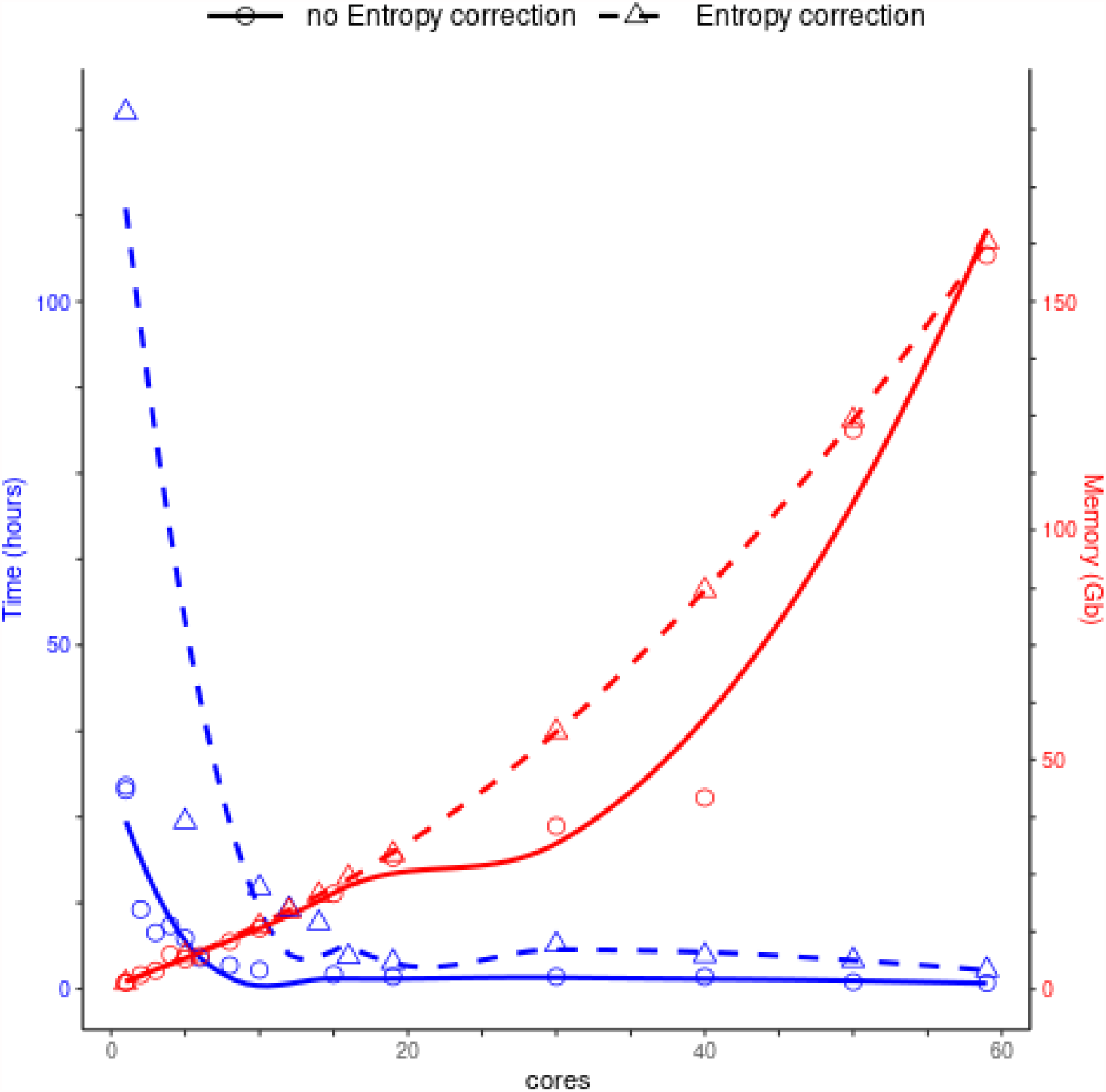
Time (blue) and memory (red) used by DnoisE to denoise and merge sequences with the Ratio-Distance criterion using different cores on a computer cluster. Denoising using entropy correction (triangles and dashed line) is compared against no correction (circles and solid line). Lines are computed using the geom_smooth() function of the ggplot2 package with method = ‘loess’.

## Conclusions

DnoisE is a novel denoising program that can be incorporated into any metabarcoding pipeline. Is it a stand alone program that addresses exclusively the denoising step, so that the users can apply their favourite programs at all other steps (e.g., chimera filtering, clustering…). Moreover, DnoisE is open-source code. Other programs used in metabarcoding pipelines also have open codes, such as DADA2 (Callahan et al., 2016), OBITOOLS (Boyer et al., 2016), SWARM (Mahé et al., 2015), or VSEARCH (Rognes et al., 2016). We strongly adhere to the open software concept for continuous and collaborative development of computing science and, in particular, in the metabarcoding field.

DnoisE is based on the UNOISE3 algorithm developed by Edgar (2016), but with three main improvements: First, it allows to select among different criteria for joining sequences; second, it incorporates the option to perform an entropy correction for coding genes; third, it is parallelizable to take advantage of the cluster architecture of modern computers.

Our correction by entropy opens a new field of analysis of coding genes, considering the different natural variability between codon positions. Antich et al (2021) found that a higher number of ESVs is retained when using entropy correction, while the proportion of erroneous sequences decreased.

In the next few years, processors are expected to reach the minimum size permitted by quantum laws. Parallel processing is needed to optimize future computer performance (Gebali, 2011; Zomaya, 2005). DnoisE offers a new parallel processing algorithm based on the MDA (maximum “daughter” abundance) to run analyses in parallel by groups of sequences that need not be compared to each other. Parallel processing allows users to run huge datasets in a fast way using multithread computers. In our example, when running with 10 cores, DnoisE took about 2.78 hours to compute a large dataset. On the other hand, memory management can be critical when running a high number of cores and large datasets and should be considered when setting the running parameters. DnoisE is written in Python3, one of the most popular languages, so it is a good option for users who want to modify or customize the code. We indeed encourage new developments of this software.

We consider that DnoisE is a good option to denoise metabarcoding sequence datasets from all kinds of markers, but specially for coding genes, given the entropy differences of codon positions. More details, sample files and complete instructions are available at github (https://github.com/adriantich/DnoisE).

## Acknowledgments

This research was funded by the the projects PopCOmics (CTM2017-88080, MICIN/AEI/FEDER,UE), MARGECH (PID2020-118550RB, MICIN/AEI/FEDER,UE), and BigPark (OAPN, 2462/2017) from the Spanish Government.

